# Dim Light at Night Disrupts the Sleep-Wake Cycle and Exacerbates Seizure Activity in *Cntnap2* Knockout Mice: Implications for Autism Spectrum Disorders

**DOI:** 10.1101/2025.03.22.644752

**Authors:** Yumeng Wang, Ketema N. Paul, Gene D. Block, Tom Deboer, Christopher S. Colwell

**Author notes:** **Correspondence:** Christopher S. Colwell. **Funding:** Supported by National Institute of Child Health Development under award number: P50HD103557 (PIs S. Bookheimer, H. Kornblum); UCLA Research Support Fund to GDB.

## Abstract

Epilepsy is one of the most common comorbidities in individuals with autism spectrum disorders (ASDs). Many patients with epilepsy as well as ASD experience disruptions in their sleep-wake cycle and exhibit daily rhythms in expression of symptoms. Chronic exposure to light at nighttime can disrupt sleep and circadian rhythms. Contactin associated protein-like 2 knockout (*Cntnap2* KO) mice, a model for autism spectrum disorder (ASD) and epilepsy, exhibit sleep and circadian disturbances and seizure-like events. This study examines how chronic dim light at night (DLaN) exposure affects sleep architecture, EEG power spectra, and seizure activity in *Cntnap2* KO and wildtype (WT) mice. Using electroencephalography (EEG) recordings, male and female *Cntnap2* KO and WT mice were exposed to DLaN (5 lux) for 2 or 6 weeks. EEG recordings were analyzed to assess sleep architecture, power spectrum, and seizure-like events. DLaN exposure delays the wake onset and disrupts sleep patterns in a sex-dependent manner, with females being more affected. DLaN significantly increased slow-wave activity (SWA, 0.5–4 Hz) in both WT and KO mice, indicating increased sleep pressure. Finally, we found that DLaN dramatically increased the frequency of seizure-like events in the Cntnap2 KO mice and even increased the occurrence rate in the WT mice. Spectral analysis of seizure-like events revealed increased theta power, suggesting the involvement of hippocampus. Chronic DLaN exposure disrupts sleep and increases seizure-like events in *Cntnap2* KO mice, with sex-specific differences. These findings emphasize the potential risks of nighttime light exposure for individuals with ASD and epilepsy, reinforcing the need to manage light exposure to improve sleep quality and reduce seizure risk.

## Introduction

Epilepsy is one of the most common comorbidities in individuals with autism spectrum disorder (ASD) [1, 2]. A systematic review and meta-analysis evaluating 66 studies, ranging from clinic samples to population samples, reported a pooled prevalence rate of epilepsy in 7% of children and 19% of adults with ASD [3]. Individuals with ASD and epilepsy tend to exhibit higher rates of intellectual disability and overall, more neurodevelopmental impairment psychiatric conditions compared with individuals with ASD without epilepsy [4–7]. Other studies have reported higher rates of epileptiform discharges and other abnormal electroencephalographic (EEG) features in individuals with ASD, even without the presence of clinical seizures. Interestingly, studies using overnight EEG monitoring in children with ASD reported higher rates of interictal epileptiform discharges (IEDs) in sleep [8–11]. These findings fit into a larger literature indicating that many patients with epilepsy experience disruptions in their sleep-wake cycle and exhibit daily rhythms in expression to symptoms [12–14]. Sleep disturbances are commonly reported in ASD, with affected individuals experiencing delayed sleep onset, nighttime arousals, fragmented sleep, and reduced total sleep duration [15–17]. The co-occurrence of epilepsy further exacerbates sleep disturbances. These associations raise questions about the underlying mechanisms, such as the shared role of pathological excitability and altered synaptic transmission that are best addressed using preclinical models [18].

To examine the intersection between autism-like behaviors, epilepsy and sleep disturbances, we searched a mouse model that exhibited all three phenotypes [19, 20]. Prior work has shown that mutations in the contactin associated protein-like 2 (*Cntnap2*) gene are associated with ASD, intellectual disabilities, and seizures in patients [21–28]. Similarly, a *Cntnap2* KO mouse model [29] has been shown to have seizures, frequent interictal discharges as well as social behavioral deficits, and repetitive behaviors [30–33]. Previous studies showed that *Cntnap2* KO mice have disrupted sleep and dampened day-night activity rhythms compared to wildtype (WT) mice [34–36]. Chronic exposure to light at nighttime has been associated with disrupted daily activity, delayed sleep onset, a dampened melatonin profile, mood alterations, metabolic dysfunctions, and poor cognition scores. We have previously shown that these mice are vulnerable to exposure to a mild, but common, circadian disruption caused by exposure to dim light at night (DLaN). This disruption impacted both activity rhythms as well as reduced social interactions and increased repetitive behaviors [36]. The negative effects of DLaN were reduced by treatment with melatonin [36] or by shifting the light toward warmer colors that minimized stimulation of melanopsin [37]. The impact of DLaN on EEG-defined sleep or epileptic discharges was not examined in the prior work.

In the present study, we used EEG/ Electromyography (EMG) recordings to measure daily rhythms in vigilance states – wake, non-rapid eye movement (NREM) sleep, and rapid eye movement (REM) sleep – in male and female *Cntnap2* KO and WT mice. Following baseline recordings under standard 12:12 light/dark conditions, the mice were exposed to DLaN for two and six weeks. We were particularly interested in evaluating the potential sex differences in the temporal patterns in vigilance states. In addition, we wanted to test the hypothesis that the *Cntnap2* KO mice would be vulnerable to circadian disruption driven by DLaN exposure. We also evaluated the impact of this environmental perturbation on the power spectrum of NREM sleep, particularly focusing on the slow-wave activity range (SWA, 0.5–4 Hz). Finally, we sought to determine if the DLaN exposure would impact the frequency of seizure-like events in the *Cntnap2* KO mice.

## Results

### Fragmented sleep and dampened daily sleep-wake rhythm in the *Cntnap2* KO mice

Prior work has found evidence for disturbed sleep/wake rhythms in male *Cntnap2* KO mice [34–36]. In the present study, we used EEG measurements to determine whether the mutation impacted daily rhythms of sleep-wake architecture in *Cntnap2* KO and WT mice. As illustrated by the hypnograms, *Cntnap2* KO mice exhibited significantly more fragmented vigilance states and increased sleep during the dark phase (**Fig. 1A**). Quantification of sleep percentages across the light and dark phases confirmed these findings, with KO mice displaying reduced sleep during the light phase and elevated sleep during the dark phase compared to WT controls (**Fig. 1B**).

**Fig 1.**
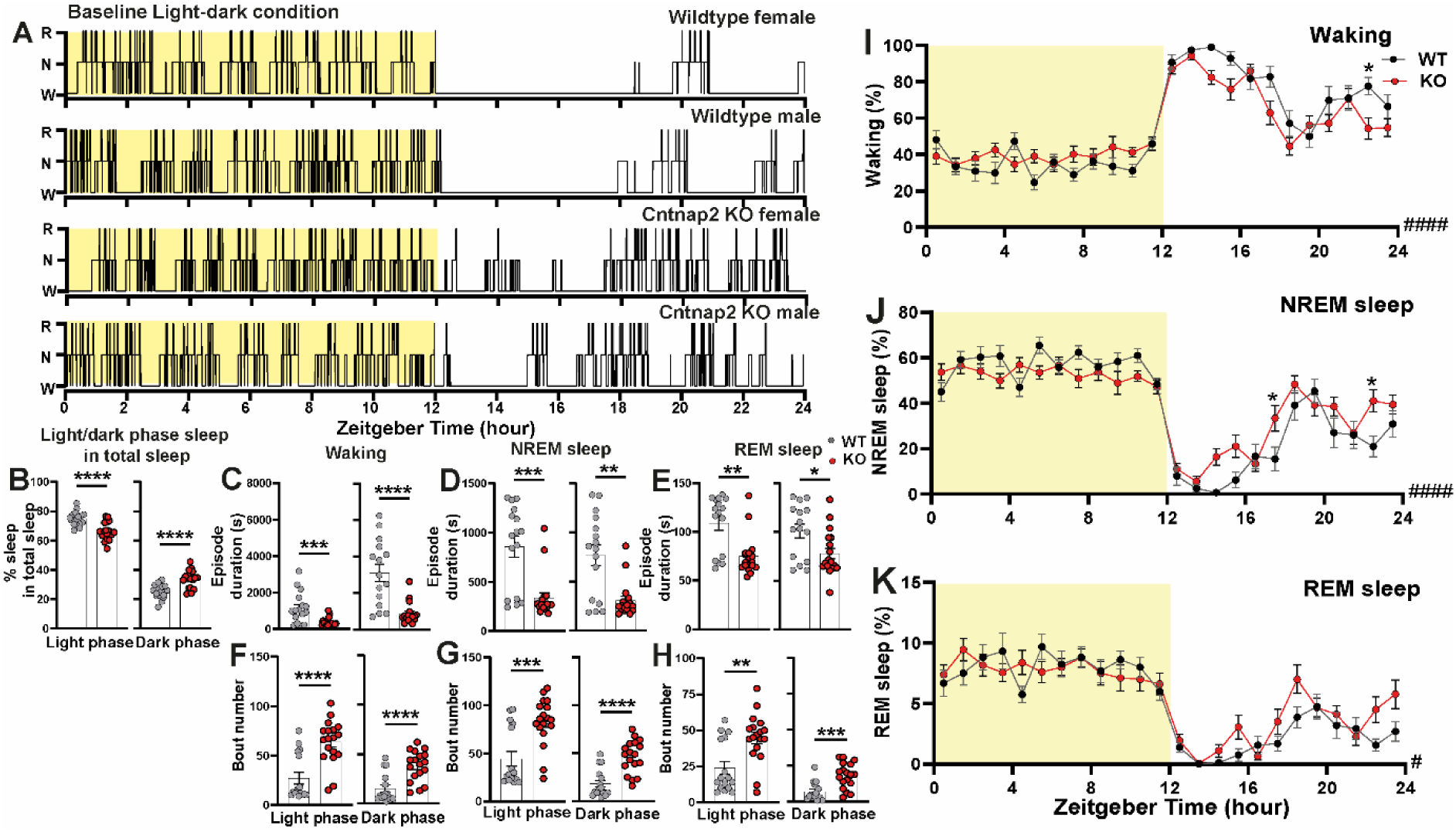
*Cntnap2* KO mice have more fragmented sleep compared with WT mice. A. Representative 24-hr hypnograms. Yellow indicates the light phase. R: REM sleep, N: NREM sleep, W: waking. B. Percentage of light and dark phase sleep in total sleep in WT (grey circle) and KO (red circle) mice. Episode duration and bout numbers of waking (C, F), NREM sleep (D, G) and REM sleep (E, H) of WT mice and KO mice under light phase (left) and dark phase (right). Asterisks indicate significant differences between WT and *Cntnap2* KO mice following either unpaired t-test or Mann-Whitney test. Time course of waking (I), NREM sleep (J), and REM sleep (K) in 1-hr bins for WT (black) and KO (red) mice. Pound sign indicates a significant interaction between the two factors “time” and “Genotype” (# p < 0.05, #### p < 0.0001, two-way ANOVA), while asterisks indicate significant differences between WT and KO mice (*p < 0.05, Bonferroni multiple comparisons test following a significant two-way ANOVA). Sample size: n = 16-18. Data are shown as mean ± SEM.

Previous studies have shown that waking episode duration is shorter in KO mice compared to WT, with a corresponding increase in waking bouts [34]. Further analysis revealed that, regardless of the light or dark phase, the average episode duration of wakefulness, NREM sleep, and REM sleep was significantly shorter in KO mice than in WT mice (**Fig. 1C-E**); similarity, the bout number for each vigilance is higher in the KO mice than in the WT mice, further supporting the presence of sleep fragmentation (**Fig. 1F-H**). A three-way ANOVA analysis of the hourly data indicated no significant difference between the sexes, or an interaction between genotype and sex, or interaction among time, genotype and sex; thus we combined the male and female data (**Table 2**). Hourly sleep-wake rhythms revealed a genotype-dependent interaction with time (**Fig. 1I-K**; **Table 1**). Post-hoc analysis indicated that KO mice slept more during the dark phase (**Fig. 1I, J**; **Table 1**). Notably, REM sleep distribution over the 24-hour cycle was affected by genotype (**Fig. 1K, Table1**).

**Table 1:** Vigilance state EEG measurements analyzed with two-way ANOVA with genotype and time as factors. For this and other tables, significant effects are shown in bold.

| Measurement | Genotype | Time | Interaction |
| --- | --- | --- | --- |
| Waking (%) | $F_{(1, 768)} = 3.828$ | $F_{(23, 768)} = 41.1$;<br>**$p < 0.0001$** | $F_{(23, 768)} = 2.813$;<br>**$p < 0.0001$** |
| NREM sleep (%) | $F_{(1, 768)} = 3.07$ | $F_{(23, 768)} = 40.43$;<br>**$p < 0.0001$** | $F_{(23, 768)} = 2.886$;<br>**$p < 0.0001$** |
| REM sleep (%) | $F_{(1, 768)} = 4.458$;<br>**$p = 0.0351$** | $F_{(23, 768)} = 23.59$;<br>**$p < 0.0001$** | $F_{(23, 768)} = 1.646$;<br>**$p = 0.0292$** |

**Table 2:** Vigilance state EEG measurements analyzed with three-way ANOVA data with sex, genotype and time as factors. Measurements were made with mice under LD 12:12 conditions and again after 6 weeks exposure to dim light at night (DLaN).

| Measurement | Time | Genotype | Sex | Time x Genotype | Time x Sex | Genotype x Sex | Time x Genotype x Sex |
| --- | --- | --- | --- | --- | --- | --- | --- |
| Waking (%) LD | $F_{(23, 216)} = 38.29$;<br>**$p < 0.0001$** | $F_{(1, 216)} = 3.903$;<br>**$p = 0.0495$** | $F_{(1, 216)} = 1.488$ | $F_{(23, 216)} = 3.009$;<br>**$p < 0.0001$** | $F_{(23, 216)} = 1.027$ | $F_{(1, 72)} = 0.051$ | $F_{(23, 72)} = 1.023$ |
| NREM sleep (%) LD | $F_{(23, 216)} = 36.38$;<br>**$p < 0.0001$** | $F_{(1, 216)} = 2.962$ | $F_{(1, 216)} = 0.851$ | $F_{(23, 216)} = 3.092$;<br>**$p < 0.0001$** | $F_{(23, 216)} = 1.001$ | $F_{(1, 72)} = 0.1469$ | $F_{(23, 72)} = 1.077$ |
| REM sleep (%) LD | $F_{(23, 216)} = 22.39$;<br>**$p < 0.0001$** | $F_{(1, 216)} = 3.905$;<br>**$p = 0.0494$** | $F_{(1, 216)} = 5.255$;<br>**$p = 0.0228$** | $F_{(23, 216)} = 1.693$;<br>**$p = 0.0288$** | $F_{(23, 216)} = 0.744$ | $F_{(1, 72)} = 0.227$ | $F_{(23, 72)} = 1.055$ |
| Waking (%) DLaN | $F_{(23, 192)} = 33.47$;<br>**$p < 0.0001$** | $F_{(1, 192)} = 1.574$ | $F_{(1, 192)} = 17.23$;<br>**$p < 0.0001$** | $F_{(23, 192)} = 2.958$;<br>**$p < 0.0001$** | $F_{(23, 192)} = 3.033$;<br>**$p < 0.0001$** | $F_{(1, 48)} = 0.713$ | $F_{(23, 48)} = 1.525$ |
| NREM sleep (%) DLaN | $F_{(23, 192)} = 30.77$;<br>**$p < 0.0001$** | $F_{(1, 192)} = 0.362$ | $F_{(1, 192)} = 14.92$;<br>**$p = 0.0002$** | $F_{(23, 192)} = 3.155$;<br>**$p < 0.0001$** | $F_{(23, 192)} = 3.034$;<br>**$p < 0.0001$** | $F_{(1, 48)} = 0.118$ | $F_{(23, 48)} = 1.417$ |
| REM sleep (%) DLaN | $F_{(23, 192)} = 26.17$;<br>**$p < 0.0001$** | $F_{(1, 192)} = 13.42$;<br>**$p = 0.0003$** | $F_{(1, 192)} = 15.22$;<br>**$p < 0.0001$** | $F_{(23, 192)} = 0.971$ | $F_{(23, 192)} = 2.423$;<br>**$p = 0.0006$** | $F_{(1, 48)} = 8.566$;<br>**$p = 0.0052$** | $F_{(23, 48)} = 1.352$ |

Further analysis of the 24-hour and light/dark phase data for each vigilance state revealed genotype- and sex-specific effects (**Table 3**). The overall time awake and REM sleep in the 24-hour period did show an effect of genotype, the KO mice had less awake time and more REM sleep compared to WT mice, and REM sleep showing a significant sex effect, males spent more time in REM sleep compared with females. During the light phase, wake time and NREM sleep were affected by the genotype, with WT mice had less wakefulness and more NREM sleep than KO mice, contributing to a blunted sleep-wake rhythm in KO mice. Additionally, light-phase wake time and REM sleep were influenced by sex, with females spending more time awake and less time in REM sleep than males. The most pronounced genotype effect was observed during the dark phase, where KO mice exhibited significantly reduced wakefulness and increased NREM and REM sleep compared to WT mice. Overall, our EEG confirmed sleep/wake data confirmed that the *Cntnap2* KO mice exhibited sleep fragmentation with a significant impact on the rhythms in wake and NREM sleep and an increase in dark-phase sleep, leading to a blunted daily sleep rhythm.

**Table 3:** Time spent in three vigilance states in WT and *Cntnap2* KO mice under LD and DLaN. Data shown as means ± SEM. a genotype difference compared in the same sex; b sex difference compared in the same genotype.

| Baseline | WT female | WT male | KO female | KO male | Genotype | Sex |
| --- | --- | --- | --- | --- | --- | --- |
| 24 hours Waking (%) | 57.82 ± 0.29 | 55.99 ± 0.34 | 54.92 ± 0.55 | 53.45 ± 0.46 | $F_{(1, 30)} = 4.458$; $p = \mathbf{0.0432}$ | $F_{(1, 30)} = 1.645$ |
| 24 hours NREM sleep (%) | 37.44 ± 0.27 | 38.83 ± 0.34 | 39.97 ± 0.54 | 40.72 ± 0.42 | $F_{(1, 30)} = 3.304$ | $F_{(1, 30)} = 0.783$ |
| 24 hours REM sleep (%) | 4.74 ± 0.05 | 5.18 ± 0.08 | 5.11 ± 0.08 | 5.82 ± 0.09 | $F_{(1, 30)} = 4.823$; $p = \mathbf{0.0359}$ | $F_{(1, 30)} = 6.146$; $p = \mathbf{0.0190}$ |
| Light phase Waking (%) | 37.72 ± 0.55 | 33.27 ± 0.75 | 41.35 ± 0.68 | 37.86 ± 0.45 | $F_{(1, 30)} = 5.446$; $p = \mathbf{0.0265}$ | $F_{(1, 30)} = 5.089$; $p = \mathbf{0.0315}$ |
| Light phase NREM sleep (%) | 54.89 ± 0.54 | 58.30 ± 0.71 | 51.65 ± 0.57 | 53.85 ± 0.41 | $F_{(1, 30)} = 5.708$; $p = \mathbf{0.0234}$ | $F_{(1, 30)} = 3.049$ |
| Light phase REM sleep (%) | 7.39 ± 0.09 | 8.43 ± 0.11 | 7.01 ± 0.18 | 8.29 ± 0.13 | $F_{(1, 30)} = 0.449$ | $F_{(1, 30)} = 8.875$; $p = \mathbf{0.0057}$ |
| Dark phase Waking (%) | 77.91 ± 0.65 | 78.70 ± 0.49 | 68.50 ± 0.80 <sup>a</sup> | 69.05 ± 0.90 <sup>a</sup> | $F_{(1, 30)} = 17.44$; $p = \mathbf{0.0002}$ | $F_{(1, 30)} = 0.087$ |
| Dark phase NREM sleep (%) | 20.00 ± 0.60 | 19.36 ± 0.39 | 28.28 ± 0.76 <sup>a</sup> | 27.59 ± 0.74 <sup>a</sup> | $F_{(1, 30)} = 17.45$; $p = \mathbf{0.0002}$ | $F_{(1, 30)} = 0.113$ |
| Dark phase REM sleep (%) | 2.09 ± 0.06 | 1.93 ± 0.13 | 3.22 ± 0.09 | 3.36 ± 0.18 | $F_{(1, 30)} = 9.502$; $p = \mathbf{0.0044}$ | $F_{(1, 30)} = 0.001$ |
| **6 weeks of DLaN** |  |  |  |  |  |  |
| 24 hours Waking (%) | 58.57 ± 0.46 | 53.41 ± 0.59 | 57.95 ± 0.26 <sup>b</sup> | 50.74 ± 0.69 | $F_{(1, 26)} = 1.340$ | $F_{(1, 26)} = 19.00$; $p = \mathbf{0.0002}$ |
| 24 hours NREM sleep (%) | 36.87 ± 0.39 | 41.71 ± 0.60 | 37.28 ± 0.24 | 42.73 ± 0.70 | $F_{(1, 26)} = 0.2690$ | $F_{(1, 26)} = 13.76$; $p = \mathbf{0.0010}$ |
| 24 hours REM sleep (%) | 4.56 ± 0.12 | 4.86 ± 0.08 | 4.77 ± 0.06 | 6.53 ± 0.26 <sup>a b</sup> | $F_{(1, 26)} = 6.789$; $p = \mathbf{0.0235}$ | $F_{(1, 26)} = 7.050$; $p = \mathbf{0.0134}$ |
| Light phase Waking (%) | 36.22 ± 0.52 | 32.71 ± 0.53 | 38.63 ± 0.50 | 35.83 ± 0.76 | $F_{(1, 26)} = 3.055$ | $F_{(1, 26)} = 3.978$ |
| Light phase NREM sleep (%) | 56.28 ± 7.50 | 59.14 ± 8.15 | 53.71 ± 0.50 | 55.06 ± 0.85 | $F_{(1, 26)} = 4.414$; $p = \mathbf{0.0455}$ | $F_{(1, 26)} = 1.770$ |
| Light phase REM sleep (%) | 7.50 ± 0.22 | 8.15 ± 0.16 | 7.66 ± 0.12 | 9.11 ± 0.25 | $F_{(1, 26)} = 1.244$ | $F_{(1, 26)} = 4.361$; $p = \mathbf{0.0467}$ |
| Dim light phase Waking (%) | 80.93 ± 0.90 | 74.11 ± 1.24 | 77.28 ± 0.42 <sup>b</sup> | 65.64 ± 1.27 | $F_{(1, 26)} = 4.804$; $p = \mathbf{0.0375}$ | $F_{(1, 26)} = 11.15$; $p = \mathbf{0.0025}$ |
| Dim light phase NREM sleep (%) | 17.45 ± 0.82 | 24.28 ± 1.13 | 20.86 ± 0.40 | 30.40 ± 1.11 | $F_{(1, 26)} = 3.661$ | $F_{(1, 26)} = 10.80$; $p = \mathbf{0.0029}$ |
| Dim light phase REM sleep (%) | 1.62 ± 0.08 | 1.57 ± 0.12 | 1.87 ± 0.05 | 3.95 ± 0.33 <sup>a b</sup> | $F_{(1, 26)} = 8.160$; $p = \mathbf{0.0083}$ | $F_{(1, 26)} = 4.901$; $p = \mathbf{0.0358}$ |

### Sleep architecture is more affected by DLaN in *Cntnap2* KO mice

To investigate the effects of DLaN on sleep architecture, we used 5 lux dim light during the dark phase, comparable to the intensity of light emitted by electronic devices in a dark room, and 300 lux during the light phase, typical of working environments. We evaluated sleep architecture in *Cntnap2* KO and WT mice over a 24-hr period following 2 and 6 weeks of DLaN exposure.

As shown in the representative hypnograms, the KO mice exhibited dramatic alterations in sleep architecture during the dim light phase after 6 weeks DLaN exposure with sex dependent effect (**Fig. 2A**). In KO females, DLaN delayed wake around the light-dark transition and increased dim light phase wakefulness (**Fig. 2H, T**, **Table 3 & 4**). Correspondingly, nighttime NREM sleep (**Fig. 2I, U**) and REM sleep (**Fig. 2J, V**) were significantly reduced (**Table 3 & 4**). In contrast, KO males showed a decrease in waking and increase NREM sleep around the light-dark transition (**Fig. 2K, L**). Notably, after just 2 weeks of DLaN exposure, KO females already exhibited reduced NREM and REM sleep during the dark phase (**Supplemental Fig. 1J-L**), indicating that changes in these vigilance states preceded the delayed wake onset observed after 6 weeks of DLaN exposure. In contrast, the WT and KO males did not show significant differences after 2 weeks of DLaN exposure (**Supplemental Fig. 1**). Although, the WT mice exhibited less pronounced changes than KO mice (**Fig. 2**), two-way ANOVA revealed significant interactions between DLaN and Zeitgeber time in WT females, indicating that DLaN altered the distribution of vigilance states across the 24-hr cycle. In contrast, WT males displayed DLaN-induced changes only in wakefulness (**Table 4**).

**Fig 2.**
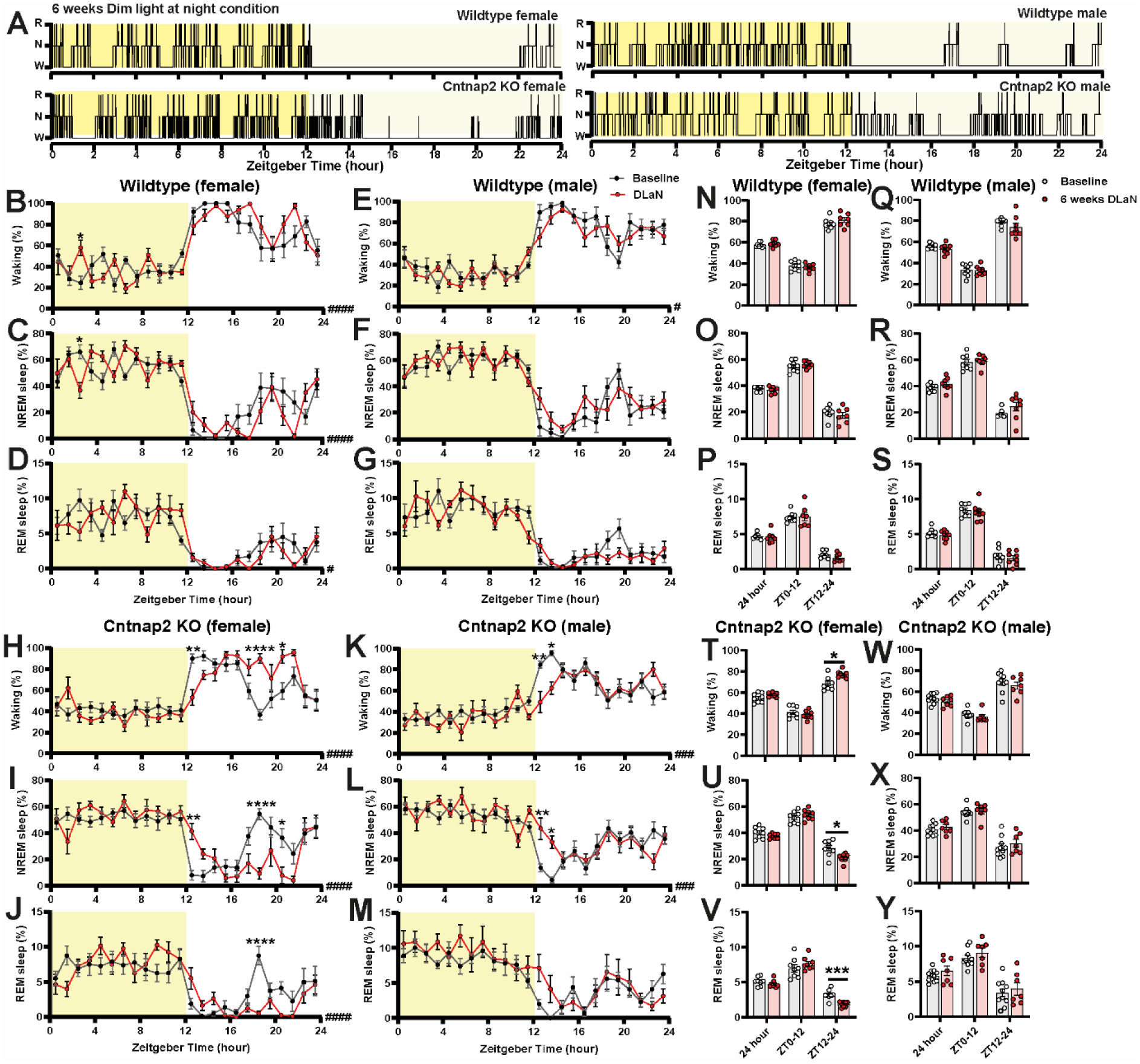
Vigilant states during baseline and 6 weeks of DLaN of WT and *Cntnap2* KO mice. A. Representative 24-hrs hypnograms. Yellow indicates the light phase and dim yellow indicates the dim light at night. (B-G, N-S) Time course of waking, NREM sleep, and REM sleep in 1-hr, 12-hr and 24-hr bins for DLaN (red) and baseline (black) conditions in wild-type female (B-D; N-P) and male (E-G; Q-S) mice. Pound sign indicates a significant interaction between the two factors “time” and “DLaN” (# p < 0.05, #### p < 0.0001, two-way ANOVA), while asterisks indicate significant differences between baseline and DLaN (*p < 0.05, Bonferroni multiple comparisons test following a significant two-way ANOVA). Sample size: n = 7–8 mice per sex per condition. (H-M; T-Y) Time course of wakefulness, NREM sleep, and REM sleep in 1-hr. 12-hr and 24-hr bins for DLaN (red) and baseline (black) conditions in *Cntnap2* KO female (H-J; T-V) and male (K-M; W-Y) mice. Pound sign from H-M indicates a significant interaction between the two factors “time” and “DLaN” (### p < 0.001, #### p < 0.0001, two-way ANOVA), while asterisks indicate significant differences between baseline and DLaN (*p < 0.05, **p < 0.01, ****p < 0.0001, Bonferroni multiple comparisons test following a significant two-way ANOVA). Asterisks in T-V indicate significant differences between baseline and DLaN (*p < 0.05, ***p < 0.001, paired t-test). Sample size: n = 7–10 mice per sex per condition. Data are shown as mean ±SEM.

**Table 4.** Two-way ANOVA for the effect of DLaN in hourly vigilance state.

| Group | Measurement | Time | Treatment | Interaction |
| --- | --- | --- | --- | --- |
| WT<br>Female | Waking (%) | $F_{(23, 312)} = 23.13;$<br>**$p < 0.0001$** | $F_{(1, 312)} = 0.1499$ | $F_{(23, 312)} = 2.904;$<br>**$p < 0.0001$** |
| | NREM sleep (%) | $F_{(23, 312)} = 23.06;$<br>**$p < 0.0001$** | $F_{(1, 312)} = 0.09997$ | $F_{(23, 312)} = 2.963;$<br>**$p < 0.0001$** |
| | REM sleep (%) | $F_{(23, 312)} = 13.27$<br>**$p < 0.0001$** | $F_{(1, 312)} = 0.1517$ | $F_{(23, 312)} = 1.779;$<br>**$p = 0.0165$** |
| WT<br>Male | Waking (%) | $F_{(23, 336)} = 25.52;$<br>**$p < 0.0001$** | $F_{(1, 336)} = 1.456$ | $F_{(23, 336)} = 1.634;$<br>**$p = 0.0348$** |
| | NREM sleep (%) | $F_{(23, 336)} = 23.80;$<br>**$p < 0.0001$** | $F_{(1, 336)} = 2.396$ | $F_{(23, 336)} = 1.539$ |
| | REM sleep (%) | $F_{(23, 336)} = 19.02;$<br>**$p < 0.0001$** | $F_{(1, 336)} = 0.8948$ | $F_{(23, 336)} = 1.448$ |
| KO<br>Female | Waking (%) | $F_{(23, 336)} = 16.03;$<br>**$p < 0.0001$** | $F_{(1, 336)} = 2.349$ | $F_{(23, 336)} = 3.819;$<br>**$p < 0.0001$** |
| | NREM sleep (%) | $F_{(23, 336)} = 15.36;$<br>**$p < 0.0001$** | $F_{(1, 336)} = 2.331$ | $F_{(23, 336)} = 3.765;$<br>**$p < 0.0001$** |
| | REM sleep (%) | $F_{(23, 336)} = 12.25;$<br>**$p < 0.0001$** | $F_{(1, 336)} = 0.8993$ | $F_{(23, 336)} = 2.908;$<br>**$p < 0.0001$** |
| KO<br>Male | Waking (%) | $F_{(23, 360)} = 14.49;$<br>**$p < 0.0001$** | $F_{(1, 360)} = 2.076$ | $F_{(23, 360)} = 2.266;$<br>**$p = 0.0009$** |
| | NREM sleep (%) | $F_{(23, 360)} = 13.28;$<br>**$p < 0.0001$** | $F_{(1, 360)} = 1.599$ | $F_{(23, 360)} = 2.277;$<br>**$p = 0.0008$** |
| | REM sleep (%) | $F_{(23, 360)} = 9.587;$<br>**$p < 0.0001$** | $F_{(1, 360)} = 3.206$ | $F_{(23, 360)} = 1.051$ |

Most previous studies on the effect of DLaN were performed in male rodents. Here, we observed distinct sex differences in sleep patterns under DLaN exposure in both WT and KO mice, particularly in total and dark/dim light-phase sleep (**Fig. 2**, **Table 3 & 4**). These findings highlight the complex interplay between sex and genotype in modulating sleep alterations under DLaN exposure, with significant effects observed across waking, NREM, and REM sleep architecture. Analysis of total 24- hr sleep revealed a significant sex difference, with KO females showed 7.2 ± 2.0% more wakefulness compared to KO males (**Table 3**). Additionally, REM sleep differed significantly across genotypes and sexes, with KO males exposed to 6 weeks of DLaN displaying a higher total amount of REM sleep than both KO females and WT males (**Table 3**). During the light phase, a significant genotypic effect was observed in NREM sleep and a sex effect in REM sleep (**Table 3**). In contrast, during the dim light phase, both genotype and sex significantly influenced sleep patterns: KO females exhibited increased wakefulness compared to KO males, while KO males had significantly more REM sleep than KO females and WT males, suggesting a sex-dependent impact of DLaN on sleep regulation (**Table 3**).

Interestingly, DLaN exposure did not affect episode durations or frequency in female mice (**Supplemental Fig. 2**). However, in male mice exposed to six weeks of DLaN, waking bout episode durations were extended relative to baseline conditions, as were NREM and REM sleep bout durations (**Supplemental Fig. 2**). These findings suggest that KO males and females may exhibit distinct phenotypes under DLaN exposure. In KO males, delayed wake onset and prolonged episode durations may serve as compensatory mechanisms to mitigate the impact of DLaN. While waking onset was also delayed in KO female, they showed increased waking time during the dim light phase, which may reduce the impact of DLaN on their sleep.

### Power spectrum of NREM sleep and the slow wave activity under DLaN

The NREM sleep power spectrum revealed a significant increase in the delta range following DLaN exposure (**Fig. 3**). In WT mice, the absolute power in the delta range of NREM sleep gradually increased over time, with a significant elevation observed after both 2 and 6 weeks of DLaN exposure compared to baseline LD conditions in the light phase (**Fig. 3A, C**, **Table 5**). Similarly, KO mice exhibited a relatively smaller but significant increase absolute power in the delta range (**Fig. 3B, D**). To further examine the spectral changes induced by DLaN, we analyzed the EEG power spectrum during waking and REM sleep, but there the changes were relatively minor. In WT mice, a small, gradual increase was observed in the theta range during REM sleep in the dark phase (**Supplemental Fig. 3B**). In KO mice, there was a slight increase in the theta range during REM sleep in the light phase of (**Supplemental Fig. 3E**).

**Fig 3.**
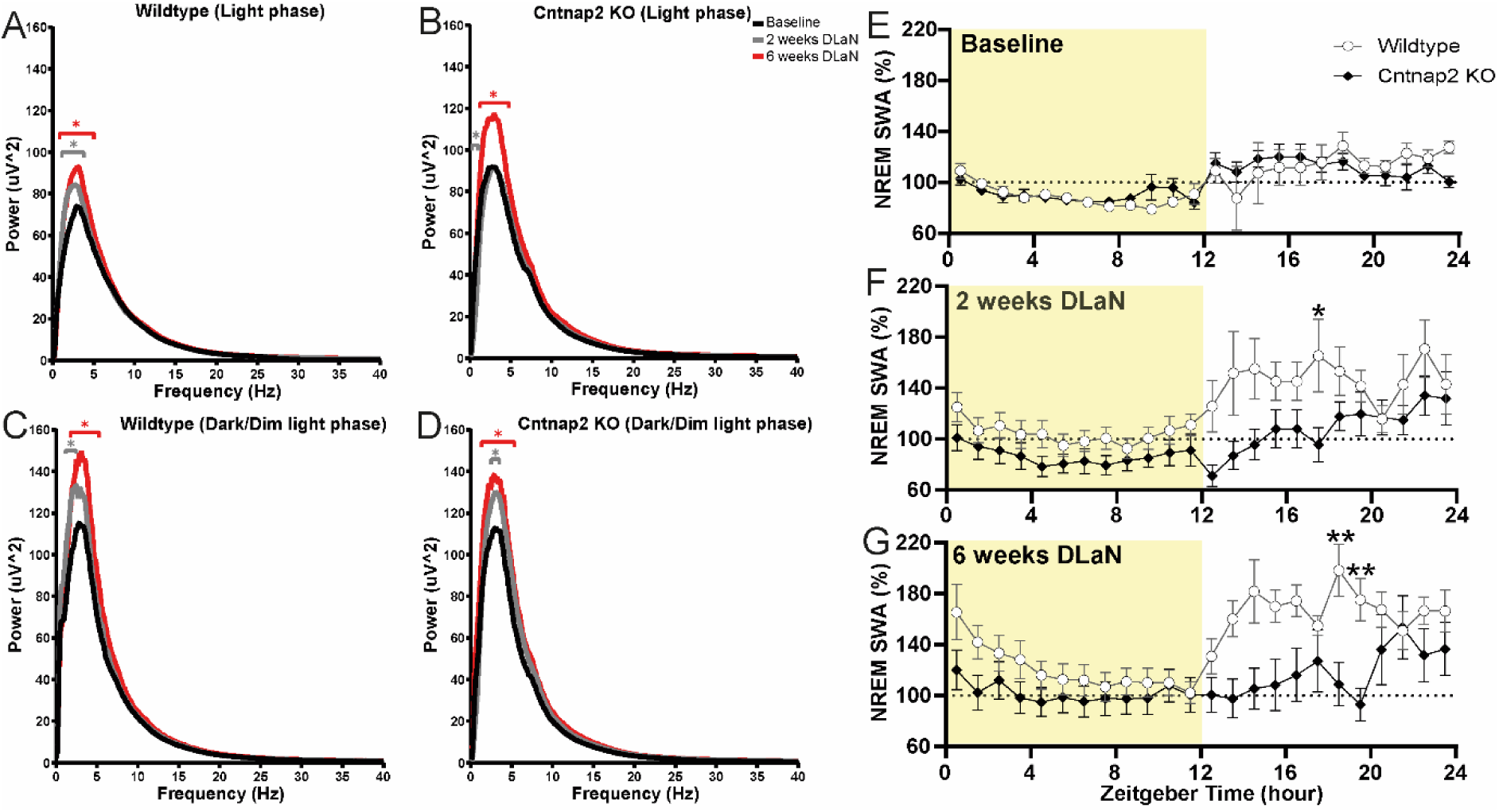
Effect of DLaN on the power spectrum and slow wave activity of NREM sleep. Absolute EEG power spectrum during NREM sleep in the light phase (A) and dark phase (C) for baseline (black), 2 weeks of DLaN (gray), and 6 weeks of DLaN (red) in wild-type mice. Asterisks indicate significant differences between baseline and DLaN (* p = 0.05–0.0001, Bonferroni multiple comparisons test following a significant two-way ANOVA for the factor “DLaN” or the interaction between “frequency” and “DLaN”). Absolute EEG power spectrum during NREM sleep in the light phase (B) and dark phase (D) for baseline (black), 2 weeks of DLaN (gray), and 6 weeks of DLaN (red) in *Cntnap2* KO mice. Asterisks indicate significant differences between baseline and DLaN (p = 0.05–0.0001, Bonferroni multiple comparisons test following a significant two-way ANOVA for the factor “DLaN” or the interaction between “frequency” and “DLaN”). The frequency bin size is 0.1 Hz. Data are shown as the mean. Time course of slow-wave activity in NREM sleep for WT (white) and *Cntnap2* KO mice (black) under baseline conditions (E), after 2 weeks (F), and after 6 weeks (G) of DLaN. Asterisks indicate significant differences between WT and *Cntnap2* KO mice (*p < 0.05, **p < 0.01; Bonferroni multiple comparisons test following a significant two-way ANOVA for the interaction of “time” and “DLaN”). Data are shown as mean ± SEM. n = 11–16 mice per genotype and condition.

**Table 5.** Power spectrum and slow wave activity in NREM sleep. Statistical results for Figure 3.

| **Group** | **Measurement** | **Frequency** | **Treatment** | **Interaction** |
| --- | --- | --- | --- | --- |
| WT light phase<br>Panel A | Power ($\mu V^2$) | $F_{(399, 14800)} = 114.5; p < 0.0001$ | $F_{(2, 14800)} = 80.43; p < 0.0001$ | $F_{(798, 14800)} = 0.908$ |
| KO light phase<br>Panel B | Power ($\mu V^2$) | $F_{(399, 16000)} = 99.3; p < 0.0001$ | $F_{(2, 16000)} = 72.07; p < 0.0001$ | $F_{(798, 16000)} = 0.732$ |
| WT DLaN<br>Panel C | Power ($\mu V^2$) | $F_{(399, 14400)} = 100.0; p < 0.0001$ | $F_{(2, 14400)} = 51.72; p < 0.0001$ | $F_{(798, 14400)} = 0.629$ |
| KO DLaN<br>Panel D | Power ($\mu V^2$) | $F_{(399, 16000)} = 112.2; p < 0.0001$ | $F_{(2, 16000)} = 66.45; p < 0.0001$ | $F_{(798, 16000)} = 0.575$ |
| **Group** | **Measurement** | **Time** | **Genotype** | **Interaction** |
| Baseline<br>Panel E | NREM SWA (%) | $F_{(23, 588)} = 8.242; p < 0.0001$ | $F_{(1, 588)} = 0.069$ | $F_{(23, 588)} = 1.269$ |
| DLaN (2 weeks)<br>Panel F | NREM SWA (%) | $F_{(23, 488)} = 4.167; p < 0.0001$ | $F_{(1, 488)} = 47.09; p < 0.0001$ | $F_{(23, 488)} = 0.752$ |
| DLaN (6 weeks)<br>Panel G | NREM SWA (%) | $F_{(23, 494)} = 3.081; p < 0.0001$ | $F_{(1, 494)} = 52.96; p < 0.0001$ | $F_{(23, 494)} = 1.249$ |

Given these observed changes in the NREM sleep power spectrum within delta range, we further analyzed the relative slow wave activity (SWA) during NREM sleep over 24 hrs, as SWA serves as a well-established marker of sleep homeostasis and sleep pressure. Under baseline LD conditions, there were no significant differences in SWA between WT and KO mice (**Fig. 3E**, **Table 5**). However, after 2 weeks of DLaN exposure, SWA increased significantly more in WT mice than in KO mice, particularly during the dark phase (**Fig. 3F**, **Table 5**). This effect became even more pronounced with extended exposure, with significant differences observed in WT mice at 6 weeks (**Fig. 3G**, **Table 5**).

### Seizure-like events increased after DLaN exposure in *Cntnap2* KO mice

During EEG scoring, abnormal bursts of high-amplitude, ‘hypersynchronized’ EEG activity were observed in *Cntnap2* KO mice during both REM sleep (**Fig. 4A**) and wakefulness (**Fig. 4B**). This abnormal EEG activity, has been previously reported in *Cntnap2* KO mice by another group and were referred to as seizure-like activity [34]. We quantified the occurrence of these seizure-like events under three conditions in both WT and KO mice. At baseline, all 16 *Cntnap2* KO mice exhibited seizure-like events (**Fig. 4C**, 100%), and these persisted under DLaN exposure. While rare in WT mice, a single seizure-like event was observed in one WT mouse at baseline. However, following 2 weeks of DLaN exposure, 6 out of 14 WT mice (42.9%) exhibited seizure-like events, increasing to 10 out of 14 WT mice (71.4%) after 6 weeks. Despite this increase in incidence, the frequency of seizure-like events in WT mice did not significantly differ between LD and DLaN conditions (**Fig. 4D**). In contrast, *Cntnap2* KO mice showed a significant increase in seizure-like events following DLaN exposure compared to baseline (Fig. 4E, mean ± SEM: baseline: 26.59 ± 5.49 events; 2 weeks DLaN: 92.47 ± 14.62 events; 6 weeks DLaN: 104.8 ± 8.43 events).

**Fig 4:**
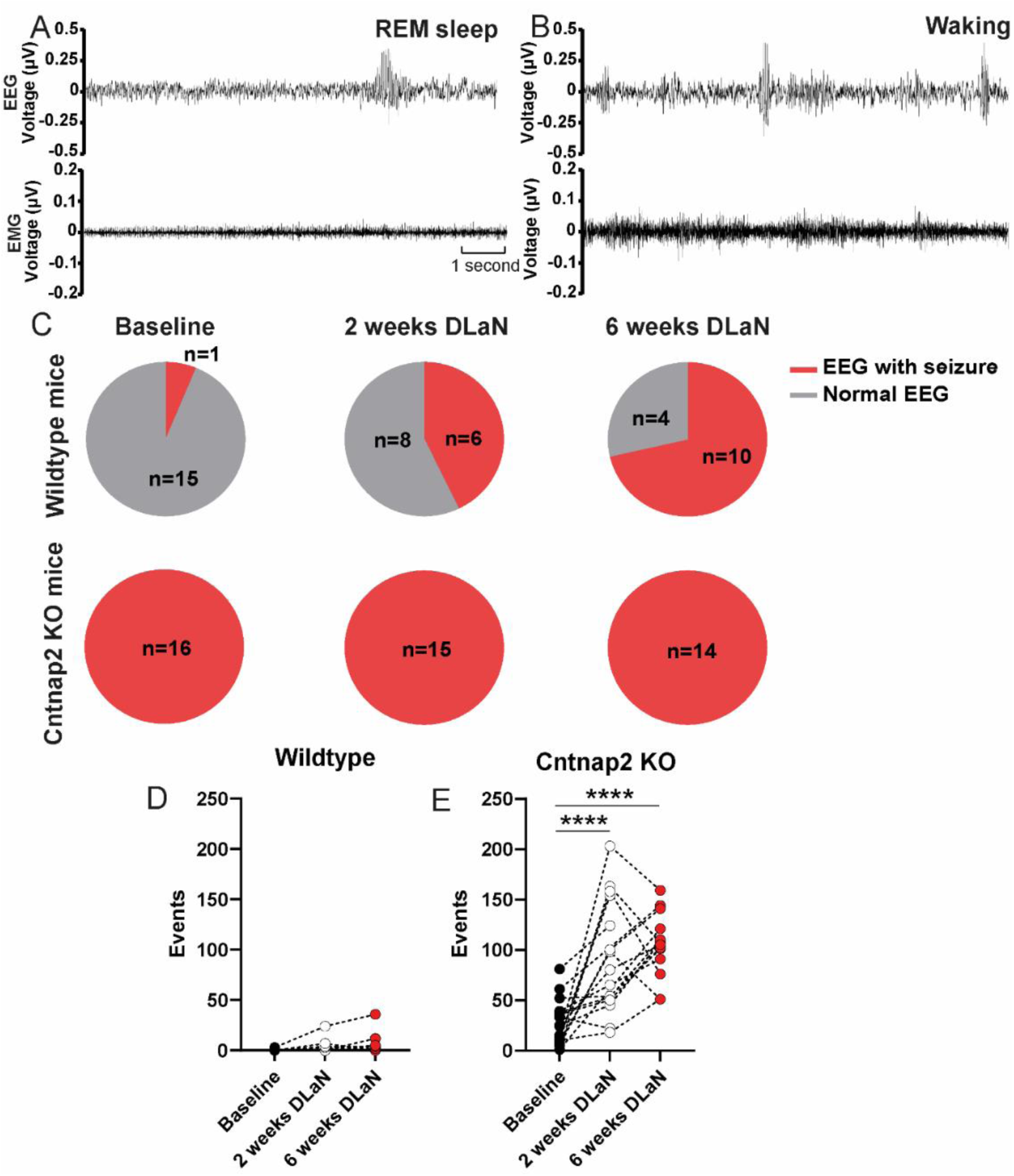
C*n*tnap2 KO mice exhibit seizure-like events. Representative events of spike-like activity during REM sleep (**A**) and waking (**B**) shown with electroencephalogram and electromyogram recordings. (**C**) Number of mice exhibiting abnormal seizure activity during baseline, after 2 weeks and 6 weeks of DLaN during 24-hr recordings of WT and *Cntnap2* KO mice. The gray portion of the pie chart with numbers indicates the number of mice showing normal EEG, while the red portion with numbers indicates the number of mice showing abnormal EEG. (**D**) Total number of events during baseline, 2 weeks and 6 weeks of DLaN in WT mice. (**E**) Total number of events during baseline, 2 weeks and of 6 weeks DLaN in *Cntnap2* KO mice. Asterisks indicate significant differences between baseline and DLaN (****p < 0.0001, Bonferroni multiple comparisons test following a significant one-way ANOVA). n = 14–16 mice per genotype and condition. Data are shown as individual values.

### The daily modulation and EEG power spectrum of the seizure-like events in *Cntnap2* KO mice

Since seizure-like events occurred during both REM sleep and wakefulness, we quantified their distribution over 24 hrs under baseline and post-DLaN conditions in KO mice. At baseline, these events were infrequent during wakefulness and occurred predominantly during REM sleep. After 2 weeks of DLaN exposure, there was a significant increase in seizure-like events during REM sleep at ZT11 (**Fig. 5A**; **Table 6**). Following 6 weeks of DLaN exposure, seizure-like events during REM sleep became more pronounced across multiple time points during the light phase (**Fig. 5C**, **Table 6**). Similarly, seizure-like events during wakefulness increased significantly after 2 weeks of DLaN exposure, particularly in the active phase (**Fig. 5B**; **Table 6**) and remained elevated after 6 weeks of DLaN (**Fig. 5D**; **Table 6**).

**Fig 5:**
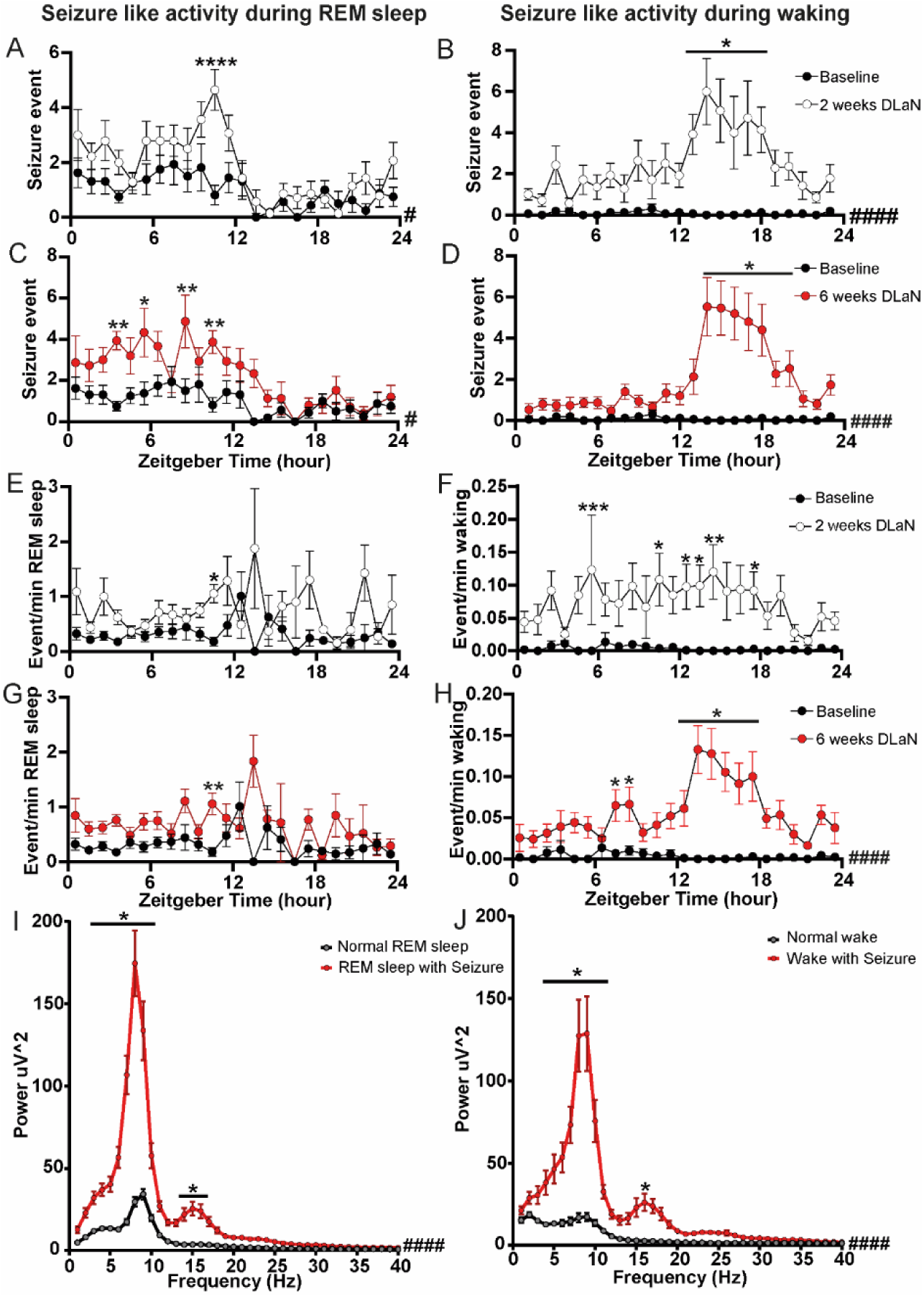
Daily distribution and power spectrum analysis of seizure-like activity in *Cntnap2* KO mice. Time course of seizure-like activity occurrence during REM sleep for *Cntnap2* KO mice under baseline (black), after 2 weeks (**A**, white), and after 6 weeks (**C**, red) of DLaN. Time course of seizure-like activity occurrence during waking for *Cntnap2* KO mice under baseline (black), after-2 weeks (**B**, white), and after 6 weeks (**D**, red) of DLaN. Asterisks indicate significant differences between baseline and DLaN (*p < 0.05, **p < 0.01, Bonferroni multiple comparisons test after significant two-way ANOVA, interaction of “time” and “DLaN”). Quantified seizure-like activity occurrence per minute of REM sleep for *Cntnap2* KO mice under baseline (black), after-2 weeks (**E**, white), and after 6 weeks (**G**, red) of DLaN. Quantified seizure-like activity occurrence per minute of waking for *Cntnap2* KO mice under baseline (black), after-2 weeks (**F**, white), and after 6 weeks (**H**, red) of DLaN. Asterisks indicate significant differences between baseline and DLaN (*p < 0.05, **p < 0.01, Bonferroni multiple comparisons test after significant two-way ANOVA, interaction of “time” and “DLaN”). Pound sign indicates a significant interaction between the two factors “time” and “DLaN” (#p < 0.05, ### p < 0.001, #### p < 0.0001, two-way ANOVA). Absolute EEG power spectrum during REM sleep (**I**) and waking (**J**) with seizure occurrence (red) and without seizure occurrence (black) for 2 weeks DLaN in *Cntnap2* KO mice. Pound sign indicates a significant interaction between the two factors “Frequency” and “DLaN” #### p < 0.0001, two-way ANOVA). Asterisks indicate significant differences between abnormal EEG and normal EEG (*p < 0.05, **p < 0.01, Bonferroni multiple comparisons test after significant two-way ANOVA, interaction of “Frequency” and “seizure”). The frequency bin size is 1 Hz. n = 12–16 mice per condition. Data are shown as mean ± SEM.

**Table 6.** Daily distribution and power spectrum of seizure-like activity. Statistical values for Fig. 5.

| Measurement | Time | Treatment | Interaction |
| --- | --- | --- | --- |
| Seizure events<br>REM sleep<br>Panel A | $F_{(23, 672)} = 5.193$;<br>$p < 0.0001$ | $F_{(1, 672)} = 35.16$;<br>$p < 0.0001$ | $F_{(23, 672)} = 1.550$; $p = 0.0489$ |
| Seizure events<br>Waking<br>Panel B | $F_{(23, 672)} = 2.710$;<br>$p < 0.0001$ | $F_{(1, 672)} = 160.6$;<br>$p < 0.0001$ | $F_{(23, 672)} = 2.848$; $p < 0.0001$ |
| Seizure events<br>REM sleep<br>Panel C | $F_{(23, 696)} = 4.457$;<br>$p < 0.0001$ | $F_{(1, 696)} = 54.81$;<br>$p < 0.0001$ | $F_{(23, 696)} = 1.707$; $p = 0.0211$ |
| Seizure events<br>Waking<br>Panel D | $F_{(23, 696)} = 5.684$;<br>$p < 0.0001$ | $F_{(1, 696)} = 169.9$;<br>$p < 0.0001$ | $F_{(23, 696)} = 6.097$; $p < 0.0001$ |
| Events/min<br>REM sleep<br>Panel E | $F_{(23, 465)} = 1.371$ | $F_{(1, 465)} = 37.06$;<br>$p < 0.0001$ | $F_{(23, 465)} = 1.548$ |
| Events/min<br>Waking<br>Panel F | $F_{(23, 624)} = 1.003$ | $F_{(1, 624)} = 139.4$;<br>$p < 0.0001$ | $F_{(23, 624)} = 1.050$ |
| Events/min<br>REM sleep<br>Panel G | $F_{(23, 500)} = 1.600$;<br>$p = 0.0390$ | $F_{(1, 500)} = 32.51$;<br>$p < 0.0001$ | $F_{(23, 500)} = 1.357$ |
| Events/min<br>Waking<br>Panel H | $F_{(23, 672)} = 3.117$;<br>$p < 0.0001$ | $F_{(1, 672)} = 204.7$;<br>$p < 0.0001$ | $F_{(23, 672)} = 3.576$; $p < 0.0001$ |
|  | **Frequency** | **Seizures** | **Interaction** |
| Power ($\mu V^2$)<br>REM sleep | $F_{(39, 1280)} = 69.92$;<br>$p < 0.0001$ | $F_{(1, 1280)} = 427.5$;<br>$p < 0.0001$ | $F_{(39, 1280)} = 31.11$; $p < 0.0001$ |
| Power ($\mu V^2$)<br>Waking | $F_{(39, 960)} = 28.69$;<br>$p < 0.0001$ | $F_{(1, 960)} = 277.0$;<br>$p < 0.0001$ | $F_{(39, 960)} = 15.58$; $p < 0.0001$ |

The observed daily rhythm in seizure-like events may be influenced by daily modulation in wakefulness and REM sleep. To account for this, we normalized seizure-like events by calculating their occurrence per minute of wakefulness or REM sleep (**Fig. 5 E-H; Table 6**). After 2 weeks of DLaN exposure, there was a significant increase in seizure-like events during both REM sleep (**Fig. 5E**; **Table 6**) and wakefulness (**Fig. 5F**; **Table 6**), with no significant interaction between time and DLaN or a main effect of time. This suggests that DLaN increased seizure-like activity without altering its temporal distribution. After 6 weeks of DLaN exposure, both DLaN and time had significant effects on seizure-like events during REM sleep, though no interaction was observed (**Fig. 5G**; **Table 6**). In contrast, seizure-like events during wakefulness showed a significant interaction between time and DLaN (**Fig. 5H**; **Table 6**), indicating the emergence of a daily rhythmic pattern after prolonged exposure.

Spectral analysis of seizure-like events after 2 weeks of DLaN exposure revealed a substantial increase in power density in the theta range, with an additional increase in the 14–16 Hz range during both REM sleep (**Fig. 5I**; **Table 6**) and wakefulness (**Fig. 5J**; **Table 6**). This increased theta power was a consistent feature across seizure-like events under both LD and DLaN conditions (**Supplemental Fig. 4**).

## Discussion

There is growing evidence that nightly exposure to light extracts a toll on our health and wellness [38, 39]. Concerns about exposure to even DLaN may be particularly relevant to individuals with neurodevelopmental disorders who are vulnerable to environmental perturbations. To explore this issue and its underlying mechanisms, we examined the impact of DLaN on the *Cntnap2* KO model of neurodevelopmental disorders. In prior work, we have shown that DLaN disrupts rhythms in activity and sleep behavior in the *Cntnap2* KO mice [36, 37]. These disruptions were correlated with DLaN-driven reductions in social behavior and increases in repetitive behaviors. While video analysis of sleep is useful for long-term, non-invasive tracking, it lacks the physiological precision of EEG, which is essential for defining sleep stages and quality. In the present study, we used EEG recordings to assess vigilance states (wake, NREM, and REM sleep) and examine the frequency of seizure-like events in the ASD model. Additionally, we explored potential sex differences, as prior work has largely focused on males.

### Sleep Disruptions in *Cntnap2* KO Mice under Baseline Conditions

*Cntnap2* is the gene that associated with both ASD and cortical dysplasia-focal epilepsy (CDFE) syndrome, and sleep problems are widely recognized in both patient populations. Sleep disruptions have been reported in other ASD mouse models, including Neuroligin-2 knockout (Nlgn2 KO) and Neuroligin-3 knockout (Nlgn3 KO) mice [40, 41], while models such as fragile X mental retardation syndrome 1 (*Fmr1*) KO and BTBR mice show genotype-time interactions rather than robust differences from WT [42]. These variations in sleep phenotypes likely stem from differences in genetic mutations leading to distinct synaptic and cellular dysfunctions [43]. Under baseline conditions, all mice exhibited robust daily rhythms in wake, NREM sleep, and REM sleep, and the vigilance states were affected by the interaction between time and genotype, only REM sleep showed a significant influence by the genotype. Prior work also found that immobility defined sleep behavior was relatively unaffected in the *Cntnap2* KO [36], but EEG-based sleep analysis revealed fragmented wakefulness and blunted state rhythms in the *Cntnap2* KO mice [34]. This is consistent with our study, comparing with the behavior defined sleep, now using EEG analysis demonstrated that sleep fragmentation is a robust feature of the *Cntnap2* KO model. Both male and female KO mice exhibited increased fragmentation of wakefulness, NREM, and REM sleep. These findings are contrast with the *Nlgn2* KO mice, which exhibit more consolidated sleep compared to WT, highlighting model-specific sleep phenotypes [41]. In the present study, both sexes of KO mice exhibited a striking increase in vigilance state fragmentation, decreased waking and increased sleep during the night, a pattern that may reflect underlying deficits in circadian regulation.

### *Cntnap2* KO mice showed dramatic changes in sleep architecture after exposure to DLaN

Under DLaN, WT mice did not show significant alterations in vigilance states after two and six weeks of exposure. These findings are consistent with a previous study demonstrating that chronic exposure to 5 lux nighttime light did not significantly alter the sleep structure of male WT mice [44]. Another study reported altered daily rhythm of vigilance state under DLaN, particularly in REM sleep [45]. In contrast to the minor changes in the WT mice, with prolonged DLaN exposure, *Cntnap2* KO mice showed disrupted sleep-wake rhythms after six weeks, indicating that the impact of DLaN increases over time. Both male and female KO mice exhibited delayed waking onset during the active phase, suggesting a similar delay effect across sexes. Delayed sleep/waking onset is similar to previous findings about the effect of the DLaN in both human and animal studies [46–48].

*Cntnap2* KO mice demonstrated increased NREM sleep irrespective of sex under baseline conditions. However, females displayed more pronounced changes in sleep architecture under DLaN, characterized by an increase in wakefulness, suggesting that females may be less susceptible to the negative effects of DLaN. Interestingly, male KO mice displayed more consolidated episodes of wakefulness, NREM sleep, and REM sleep under DLaN conditions. In contrast, female KO mice and WT mice did not exhibit significant changes in sleep architecture under these conditions. These sex-specific responses may be driven by hormonal differences. Given that ASD is predominantly studied in males [49], our inclusion of both sexes revealed subtle but significant sex differences in response to DLaN. Similar sex-specific responses to nighttime light have been observed in other species, such as birds, where females exhibit greater wakefulness under nocturnal light exposure [50]. These differences highlight the potential influence of environmental factors and hormonal regulation on the sleep phenotypes of males and females under DLaN conditions.

Spectral analysis of the EEG during NREM sleep revealed a significant increase in delta power in both WT and KO mice under DLaN compared to baseline LD conditions, indicating elevated sleep pressure. This finding aligns with prior studies reporting similar effects after chronic DLaN exposure for durations ranging from one week to one month [46]. A 24-hr profile of slow-wave activity (SWA) data demonstrated more pronounced changes during the dark phase in WT mice. Although KO mice exhibited increased delta power in their NREM sleep power spectrum, these mice also had increased sleep during the dim light phase that may result in decreased relative SWA. In contrast, the extended waking periods in WT mice likely contributed to their elevated SWA. Both spectral data and relative SWA data suggest that KO and WT mice experienced greater sleepiness under DLaN conditions compared to baseline LD conditions.

### Seizure-like activity occurrence increased under DLaN conditions

Epileptic seizures and abnormal EEG patterns have been previously observed in *Cntnap2* KO mice [30, 34]. Consistent with previous findings, we observed abnormal electrical discharges in both male and female KO mice, indicating that this EEG feature is present across sexes. Under baseline conditions these seizure-like events occurred primarily during REM sleep and were rarely observed during waking and never during NREM sleep. While, in general, approximately half of all seizures occur during sleep, most sleep-related seizures are associated with NREM sleep, with REM-related seizures being relatively rare [51]. Similarly, *Ank3-exon1b* knockout (Ank3-1b KO) mice, a model relevant to epilepsy, also exhibited a higher prevalence of seizures during REM sleep [52], and *Nlgn2* KO mice also showed seizure-like events predominated during wakefulness and REM sleep [53]. These findings suggest that abnormal EEG activity in epilepsy and autism models is strongly associated with REM sleep.

As has been reported in patients and rodents, seizures are likely to occur at specific times, and the circadian rhythm of seizure occurrence varies among different epilepsy subtypes and location of seizure foci [18, 54, 55]. In *Nlgn2* KO mice, hypersynchronized ECoG events peaked around ZT 9– 10, while in *Cntnap2* KO mice, seizure density was highest at the beginning of the active phase under LD conditions. Our study found that chronic DLaN exposure significantly increased seizure-like event frequency in KO mice in REM sleep and waking. After six weeks of DLaN, seizure density exhibited a distinct rhythmic pattern in waking, suggesting that prolonged exposure to dim light may trigger adaptive mechanisms. Notably, WT mice also displayed seizure-like events under DLaN, though at a lower frequency, indicating that DLaN may be an environmental risk factor for both KO and WT mice.

### Excitatory/inhibitory imbalance in *Cntnap2* KO mice

Prior work suggests that network-wide differences from WT might underlie the increased occurrence of seizure-like events in *Cntnap2* KO mice. *Cntnap2* expression is highest in cortical and striatal regions [56], and its loss leads to dysregulated inhibitory synaptic transmission, and impaired neuronal migration. Power spectral analysis revealed that these seizure-like events were characterized by elevated theta power, which is likely to be generated by the hippocampus suggesting that the hippocampus may play a critical role in the generation of these seizure-like events. A reduction in parvalbumin-positive interneurons in the hippocampus has been described in *Cntnap2* KO mice [30]. Dysfunction of parvalbumin-positive (PV+) interneurons has been implicated in several neurological disorders, including epilepsy [57, 58]. PV interneuron failure, potentially via depolarization block and impaired inhibitory synaptic function, results in a reduction in overall inhibition and ultimately in seizures or seizure-like events in the *Cntnap2* KO mice.

In addition to PV neuron dysfunction, *Cntnap2* KO animals have been reported to exhibit reduced functional synaptic connectivity and decreased synapse density in the medial prefrontal cortex [59, 60]. Notably, modulation of the excitatory/inhibitory (E/I) balance in the prefrontal cortex has been shown to rescue social behavior deficits in *Cntnap2* KO mice [61]. A recent study demonstrated that the daily E/I balance was disrupted in *Fmr1* KO and BTBR mice, two other autism mouse models [42], suggesting that disrupted daily E/I balance may be a common feature of autism-related models. Environmental factors, such as DLaN, may further exacerbate these network imbalances. Chronic exposure to 5 lux DLaN has been shown to alter thalamo-cortical neuronal networks in animal studies [46]. In humans, sleeping under a night light of approximately 40 lux has been associated with shallower sleep, increased arousals, and significantly decreased brain oscillations during sleep [62]. Given that DLaN can disrupt brain networks, and *Cntnap2* KO mice already exhibit disrupted E/I balance, exposure to DLaN likely intensifies these pre-existing neural deficits. This compounded disruption may explain the increased occurrence of seizure-like events observed in KO mice under DLaN conditions, as compared to the baseline LD condition.

### Disrupted circadian rhythm is a risk factor for seizure occurrence in *Cntnap2* KO mouse model

The circadian clock drives daily rhythms in physiology through a variety of mechanisms. In the mouse cortex, the majority of synaptic transcripts exhibit daily rhythms in abundance that peak before dawn and dusk [63] and robust rhythms in phosphoproteins have been described [64]. In prior work, we have shown that DLaN altered the phase and amplitude of the molecular clock expressed in the *Cntnap2* KO in a tissue-specific manner [36] and is likely to alter cortical physiology. Previously an interaction between disruption of the circadian clock and epilepsy has been suggested. For example, mutations in the RORα gene (RORA) link to intellectual disability including autistic symptoms [65]. In another example, mutations in SCN1A which encodes a voltage-gated sodium channel (Nav1.1), which is necessary for normal circadian rhythms, and is a risk factor for epilepsy [66, 67]. Similarly, a Nav1.1 mutation causing reduced interneuron excitability and seizures also causes sleep impairment in a mouse model of Dravet Syndrome [68].

The evidence for bi-directional associations between circadian disruption and susceptibility to epilepsy are intriguing. DLaN is a mild environmental disruptor of circadian rhythms and one that may be commonly experienced by patients with ASD and other neurodevelopmental disorders. Our work raises the possibility that sleep and circadian disruption by environmental factors makes at least some of the core symptoms of developmental disabilities worse. Our findings also raise the possibility of using circadian medicine approaches to develop new interventions. Many have argued for the possible benefits of interventions at specific times of the day to strengthen the patients’ circadian rhythms. In this mouse model, we have already shown that nighttime treatment with melatonin improves the circadian rhythms as well as autistic-like behavior [36]. The same treatment during the day did not produce these benefits. The timed treatment of melatonin supplements or melatonin receptor agonists may prove to be an appropriate countermeasure. It will be important to examine whether this class of drugs may also be effective in reducing the epileptic discharges. Here it is worth noting that the epilepsy associated with focal cortical dysplasia and *Cntnap2* mutations is commonly resistant to pharmacological management and new treatments are badly needed.

### Limitations

There are several limitations to this work that will need to be addressed in future studies. This study exclusively uses *Cntnap2* KO mice while ASD and epilepsy are genetically heterogeneous, and results from one model may not generalize to other forms of these disorders. Future research should assess whether the effects observed here are consistent across other ASD-related mouse models following DLaN exposure.

Additionally, while our findings show that DLaN increases seizure susceptibility, the mechanism of action remains unclear. Specifically, it is not clear whether the effects of DLaN are due to circadian rhythm disruption, direct light exposure, or a combination. To address this, future studies should manipulate the circadian system independently of DLaN exposure to determine whether the observed effects are driven specifically by circadian mechanisms.

## Conclusion

Our study highlights the significant negative impact of DLaN on sleep and neurological health, underscoring the importance of addressing this increasingly common environmental factor. Unlike many previous studies focusing on male rodents, we included both sexes and found that female mice were less vulnerable to DLaN-induced waking disruptions compared to males. *Cntnap2* KO were particularly sensitive to DLaN, showing more significant sleep disruptions and a sex-dependent effect on sleep phenotype.

A striking finding was the increased occurrence of seizure-like activity under prolonged DLaN exposure. While *Cntnap2* KO mice naturally displayed seizure-like activity due to their genetic deficit, we observed an increase in the occurrence under DLaN conditions, not only in *Cntnap2* KO mice, but also in the WT mice. This suggests that chronic DLaN exposure can alter brain networks, potentially exacerbating neurological vulnerabilities. Our results emphasize the urgent need to optimize lighting environments, particularly for individuals with ASD and epilepsy, who may be more sensitive to the effects of artificial light at night. Promoting sleep hygiene, such as minimizing exposure to dim light before and during sleep, could mitigate these risks. These findings not only extend our understanding of DLaN’s impact on vulnerable populations but also serve as a call to action for improving lighting habits in daily life to safeguard both neurological and overall health.

### Methods Animals

Mice (13-15 weeks old) were used in this study. All animal procedures were performed in accordance with the UCLA animal care committee’s regulations. *Cntnap2*^tm2Pele^ mutant mice backcrossed to the C57BL/6J background strain were acquired (B6.129(Cg)-*Cntnap2*^tm2Pele^/J, https://www.jax.org/strain/017482 [29]. *Cntnap2* null mutant (KO) and C57BL/6J wild-type (WT) mice were obtained from heterozygous breeding pairs. Weaned mice were genotyped (TransnetYX, Cordova, TN) and group-housed prior to experimentation. Mice were housed in light-tight ventilated cabinets in temperature- and humidity-controlled conditions, with free access to food and water. Male and female mice will be used in this study.

### EEG/EMG Surgery

At the age of 13-15 weeks, animals were operated under Ketamine/Xylazine anesthesia (100/8.8 mg/kg, intraperitoneal injection) and EEG/ EMG electrodes were implanted as described previously [69]. A prefabricated head mount (CAT:8201, Pinnacle Technologies, KS) was used to position four stainless-steel epidural screw electrodes. One front electrode was placed 1.5 mm anterior to bregma and 1.5 mm lateral to the central line, whereas the second two electrodes (interparietal—located over the visual cortex and common reference) were placed 2.5 mm posterior to bregma and 1.5 mm on either side. A fourth screw served as a ground. Electrical continuity between the screw electrode and head mount was aided by silver epoxy. EMG activity was monitored using stainless-steel Teflon-coated wires that were inserted bilaterally into the nuchal muscle. The head mount (integrated 2 × 3 pin grid array) was secured to the skull with dental cement. Mice were allowed to recover for at least 7 days before sleep recording.

### Experimental design

*Cntnap2* KO littermate and age-matched WT littermate controls were first entrained to a normal lighting cycle: 12 h light: 12 h dark (LD). Light intensity during the day was 300 lx as measured at the base of the cage, and 0 lx during the night. One week after surgery, mice were connected to a lightweight tether attached to a low-resistance commutator mounted over the cage (Pinnacle Technologies, KS) as previously described [69]. Mice were allowed a minimum of 2 additional days to acclimate to the tether and recording chamber. Subsequently, a baseline (BL) day was recorded, starting at lights on zeitgeber time 0. After the baseline recording, the mice were housed under dim light at night (DLaN; ZT0-12: 300 lux; ZT12-24: 5 lx) for 2 weeks as described previously [36]. Two weeks later, the mice were attached to the commutator for the post 2 weeks DLaN recording for 24 hrs. After the recording, the mice were continuously housed under DLaN for another 4 weeks. After a total 6 weeks of DLaN, the mice were again attached to the commutator for the post 6 weeks DLaN recording for 24 hrs.

### EEG data acquisition and EEG power spectrum analysis

Data acquisition was done by Sirenia Acquisition software (Pinnacle Technologies, KS) as previously described [69]. EEG signals were low pass filtered with a 40 Hz cutoff and collected continuously at a sampling rate of 400 Hz. All data were recorded simultaneously in 1s epoch. Waking, NREM, and REM sleep were determined, movement artifacts and abnormal EEG events were excluded for power spectral and slow-wave analysis. For the spectral analysis, complete and clean recordings are needed to enter the analysis. Unfortunately, this was not possible in six animals (four male KO and two male WT mice), which were therefore excluded from the analysis of the absolute EEG power density spectra and SWA in NREM sleep. Spectral analysis was performed using fast Fourier transform (FFT; 0.1–40 Hz, 0.1 Hz resolution); the absolute power density spectra of waking and REM sleep in baseline and post DLaN days were analyzed.

### Data analysis

Three vigilance states (waking, NREM, and REM sleep) were scored offline in 10-s epochs by the Sirenia program. The manual scoring of vigilance states based on the EEG and EMG recordings was performed according to standardized criteria for mice [46]. The average amount of the vigilance states (waking, NREM sleep, and REM sleep) and EEG spectral data and NREM sleep SWA were analyzed in 1 hr, 12 hr and 24 hr intervals.

Episode duration averages were compiled by the Sirenia program, which uses a conservative algorithm for bout lengths requiring sustained changes in arousal state to switch state. A bout was defined as 3+ continuous epochs, and the occurrence of 3+ continuous epochs of the different stage indicated the end of the current bout. Episode duration and bout number were analyzed in 12 hr intervals.

To investigate the effect of DLaN on EEG power density of NREM sleep, we analyzed the relative EEG power density in the slow-wave range (SWA, 0.5–4.0 Hz) in NREM sleep as described previously [46]. Since for the slow wave activity analysis, it was necessary to re-calculate power density values relative to the control condition (WT, n = 14; KO, n = 17), complete and clean recordings were needed from all animals for both conditions to enter the analysis. Unfortunately, this was not possible in 5 animal under 2 weeks of DLaN and 5 of 6 weeks DLaN, which were therefore excluded from the analysis of SWA (2 weeks DLAN: WT, n = 11, KO, n = 15; 6 weeks DLaN: WT, n = 12, KO, n = 14).

### Seizure-like events identification

During vigilance state identification, occasional and distinct events of high amplitude EEG bursts were observed. Since alteration in E/I balance has been postulated as a mechanism underlying epileptogenesis and seizure generation, this abnormal EEG activity might be indicative of hypersynchronisation and/or epileptiform activity. Thus, these abnormal EEG events were marked on EEG traces (and excluded from the spectral analysis of the vigilance states described above). More precisely, the number and duration of abnormal events were quantified by marking them according to the following three criteria under either waking or REM sleep [41]: amplitude at least twice that of the background EEG signal, duration of at least one second and not caused by movement of the animal (no movement artifacts). Two events separated by less than 0.5 s were considered as a single one.

### Seizure-like activity events occurrence and power spectrum analysis

To evaluate seizure-like events, we quantified their occurrence in 1-hr and 24-hr intervals. Given the differences in the amount of waking and REM sleep across the 24-hr cycle, we further normalized the frequency of these EEG events, calculating them as events per minute of REM sleep or waking under baseline conditions, after 2 weeks, and after 6 weeks of DLaN.

To better capture these EEG events, we score the EEG data in 1 second epoch during both normal REM sleep or waking states and during periods exhibiting seizure-like activity. EEG power density was calculated within the 1–40 Hz range (1-Hz resolution) for these abnormal events and compared to the power density during normal waking or REM sleep occurring immediately before or after the seizure-like events.

### Statistical analysis

For data analysis, GraphPad Prism 10 (GraphPad software) was used. A two-way analysis of variance (ANOVA) was used to compare the effect of DLaN across time, or EEG frequencies. If the result was significant, a post hoc Bonferroni multiple comparisons corrected t test was performed, and the significant time points and EEG frequencies bins are reported. Paired student’s t-tests were used when distributions were normal; otherwise, the nonparametric Wilcoxon matched-pairs signed rank test was used to compare the difference between baseline and DLaN conditions.

Given the primary focus on the effects of DLaN, most comparisons were made between the LD and DLaN conditions. Unpaired student’s t-tests were used when distributions were normal; otherwise, nonparametric Mann-Whitney tests were used to determine statistically significant differences between WT and *Cntnap2* KO mice. To account for potential confounding factors such as genotype and sex, a three-way ANOVA was conducted under three distinct conditions. The detailed data, including associated F-statistics, are presented in Table 2. A two-way ANOVA was used to compared the difference the genotype and sex difference under light, dark/dim light phase and total 24-hr under either LD or DLaN conditions, if the result was significant, a post hoc Bonferroni multiple comparisons corrected t test was performed, details are presented in Table 3. Data are reported as mean and standard error of the mean (SEM), and the threshold for statistical significance was set to 0.05.

## Abbreviations

Ank3-1b: Ank3-exon1b
ANOVA: analysis of variance
ASD: autism spectrum disorder
BL: baseline
CDFE: cortical dysplasia-focal epilepsy
Cntnap2: contactin associated protein-like 2
DLaN: dim light at night
EEG: electroencephalograph
EMG: electromyographic
E/I: excitatory/inhibitory
Fmr1: fragile X mental retardation syndrome 1
Nlgn: neuroligin
NREM: non rapid eye movement
n.s.: not significant
PV+: parvalbumin-positive
KO: knockout
LD: light-dark
SWA: slow-wave activity
WT: wild type
ZT: Zeitgeber time

## Acknowledgments

We extend our gratitude to Drs. Cristina A. Ghiani and Kathy Tamai for the useful discussions throughout the study. We are also grateful to Dr. Keith Lewy, the veterinarian from the animal facility, for providing post-surgical care for the mice.

## Author contributions

CSC and YW conceptualized the experiments. YW conducted the experiments and analyzed the data. YW, TD and CSC identified the seizure-like events. YW, TD and CSC wrote and manuscript. All authors read and approved of the final manuscript.

### Funding

Supported by National Institute of Child Health Development under award number: U54HD087101 (PIs S. Bookheimer, H. Kornblum); UCLA Research Support Fund to GDB

## Data availability

The datasets here used and analyzed are available from the corresponding author upon request.

## Declarations Ethical approval

All animal procedures were performed in accordance with the UCLA animal care committee’s regulations.

## Supplemental Figures

**Supplemental Fig 1:**
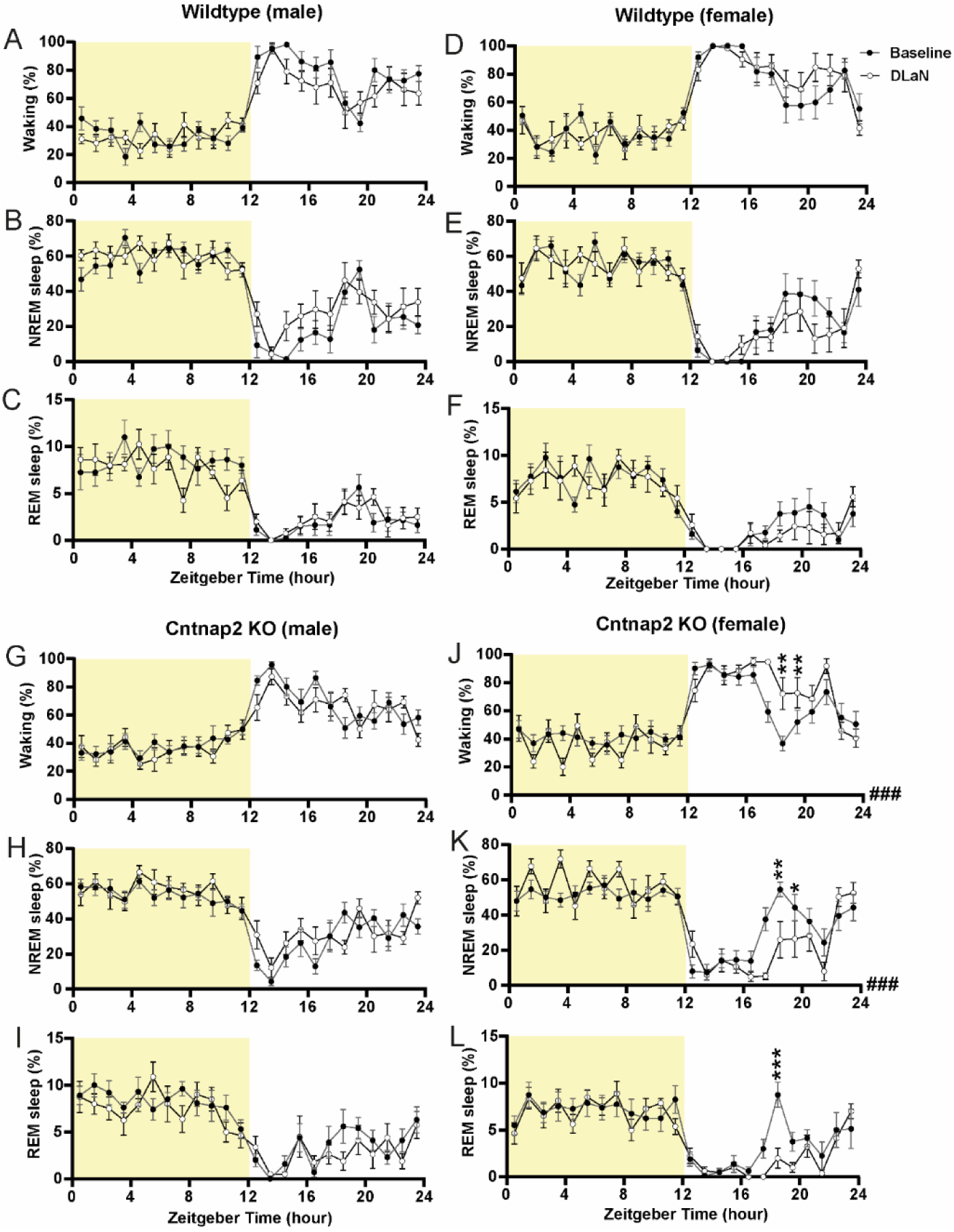
Vigilant states during baseline and 2 weeks of DLaN in WT and *Cntnap2* KO mice. **(A–F)** Time course of waking, NREM sleep, and REM sleep in 1-hour bins for DLaN (white) and baseline (black) conditions in wild-type male (**A, B, C**) and female (**D, E, F**) mice. Sample size: n = 7–8 mice per sex per condition. **(G–L)** Time course of wakefulness, NREM sleep, and REM sleep in 1-hr bins for DLaN (white) and baseline (black) conditions in *Cntnap2* KO male (**G, H, I**) and female (**J, K, L**) mice. Pound sign indicates a significant interaction between the two factors “Zeitgeber time” and “DLaN” (### p < 0.001, two-way ANOVA), while asterisks indicate significant differences between baseline and DLaN (*p < 0.05, **p < 0.01, ***p < 0.001, Bonferroni multiple comparisons test following a significant two-way ANOVA). Sample size: n = 7–10 mice per sex per condition. Data are shown as mean ± SEM.

**Supplemental Fig 2:**
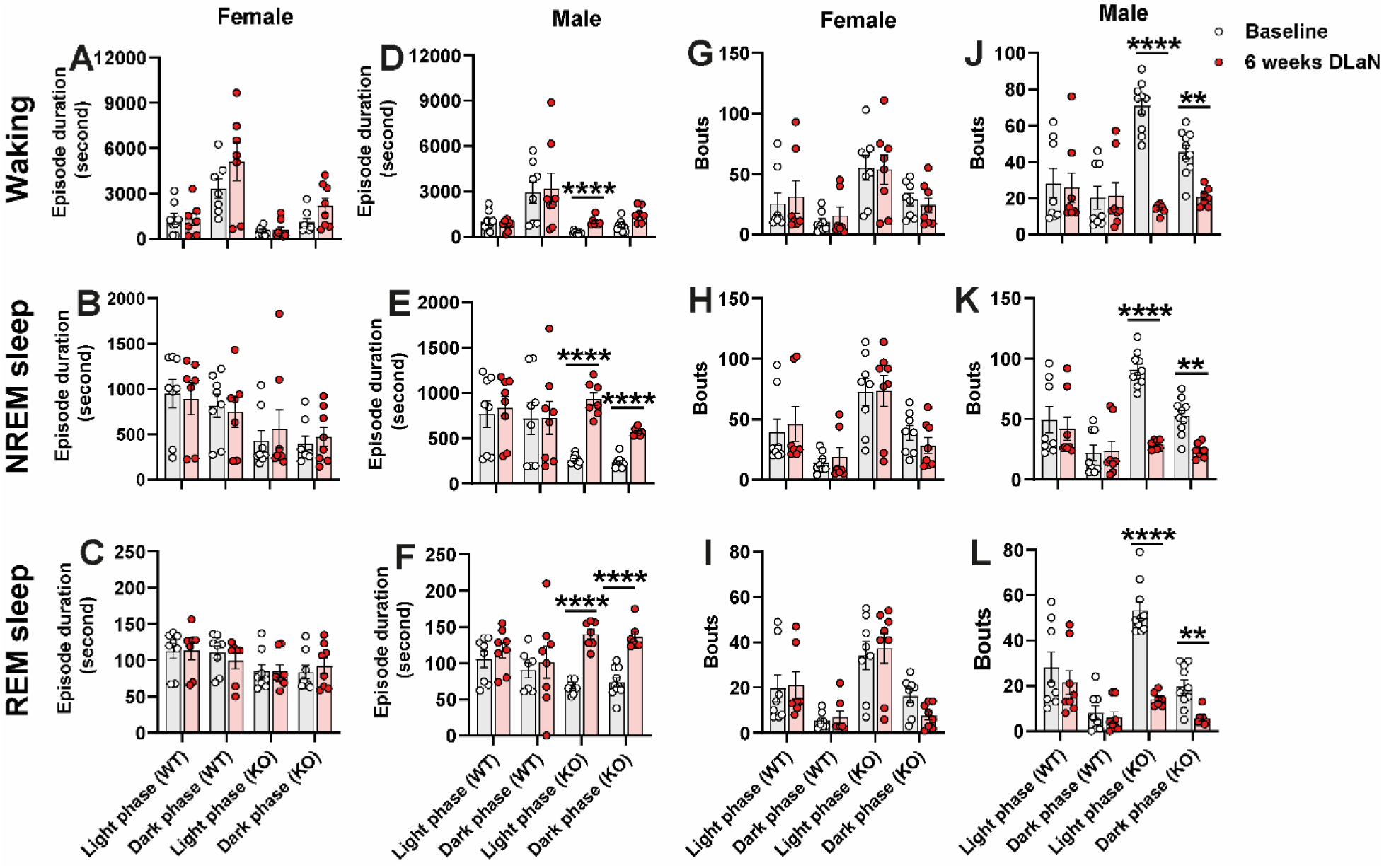
Episode duration and bout number in baseline and DLaN in WT and *Cntnap2* KO mice. (A–F) Episode duration of waking, NREM sleep, and REM sleep for baseline (white circle) and 6 weeks DLaN (red circle) in female (A, B, C) and male (D, E, F) mice. (G-L) Bout number of waking, NREM sleep, and REM sleep for baseline (white circle) and 6 weeks DLaN (red circle) in female (A, B, C) and male (D, E, F) mice. Asterisks indicate significant differences between baseline and DLaN (*p < 0.05, **p < 0.01, ***p < 0.001, ****p < 0.0001, paired t -test or Wilcoxon matched-pairs signed rank test). Sample size: n = 7–10 mice per sex, genotype, and condition. Data are shown as mean ± SEM.

**Supplemental Fig 3:**
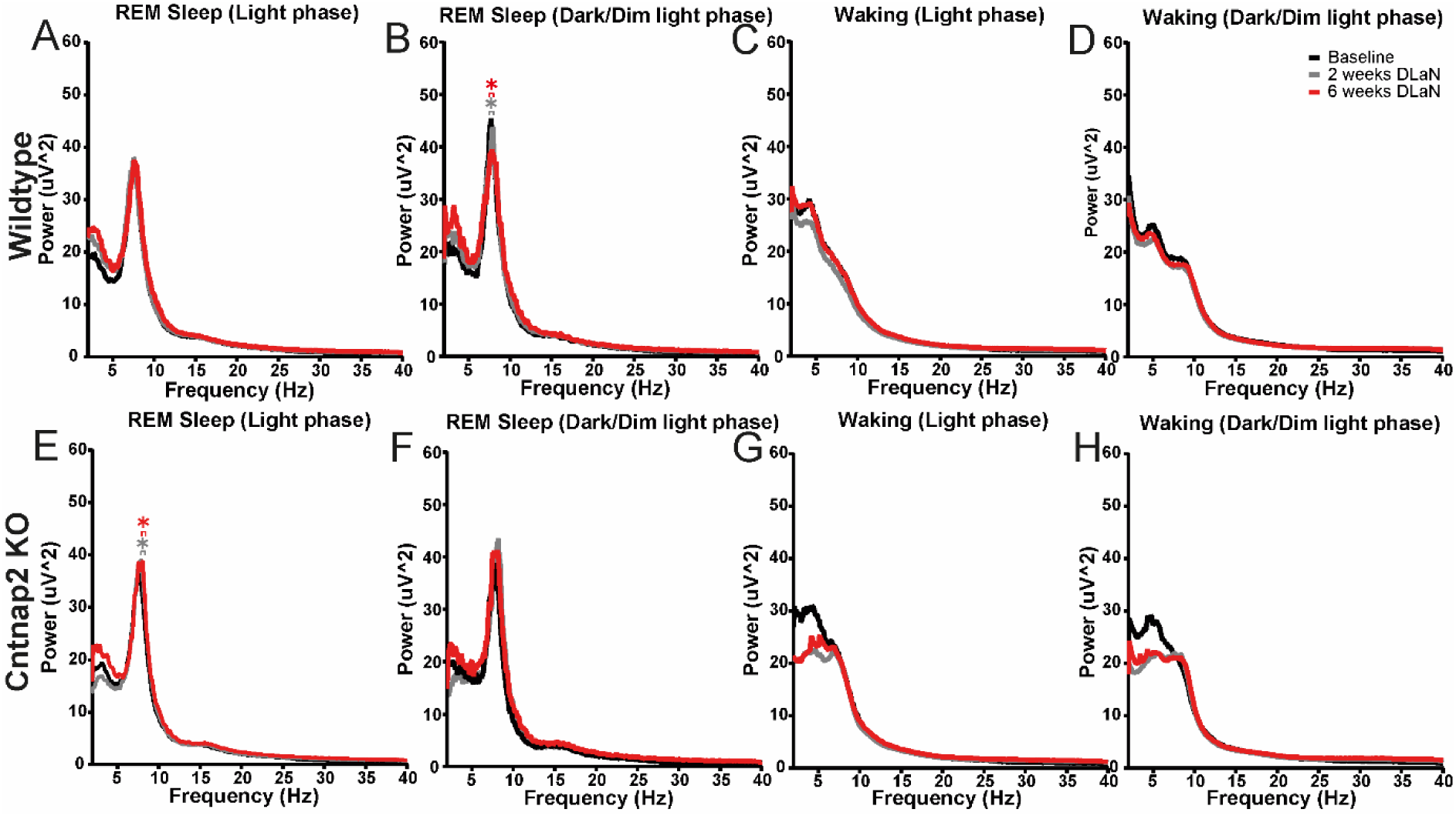
Effect of DLaN on the power spectrum of REM sleep and waking. Absolute EEG power spectrum during REM sleep in the light phase (**A**) and dark/dim light phase (**B**) for baseline (black), 2 weeks of DLaN (gray), and 6 weeks of DLaN (red) in wild-type mice. Asterisks indicate significant differences between baseline and DLaN (p = 0.05–0.0001, Bonferroni multiple comparisons test following a significant two-way ANOVA for the factor “DLaN”). Absolute EEG power spectrum during waking in the light phase (**C**) and dark/dim light phase (**D**) for baseline (black), 2 weeks of DLaN (gray), and 6 weeks of DLaN (red) in wild-type mice. Absolute EEG power spectrum during REM sleep in the light phase (**E**) and dark/dim light phase (**F**) for baseline (black), 2 weeks of DLaN (gray), and 6 weeks of DLaN (red) in *Cntnap2* KO mice. Absolute EEG power spectrum during waking in the light phase (**G**) and dark/dim light phase (**H**) for baseline (black), 2 weeks of DLaN (gray), and 6 weeks of DLaN (red) in *Cntnap2* KO mice. The frequency bin size is 0.1 Hz and X axis started from 3 Hz. n = 11–16 mice per genotype and condition. Data are shown as the mean.

**Supplemental Fig 4:**
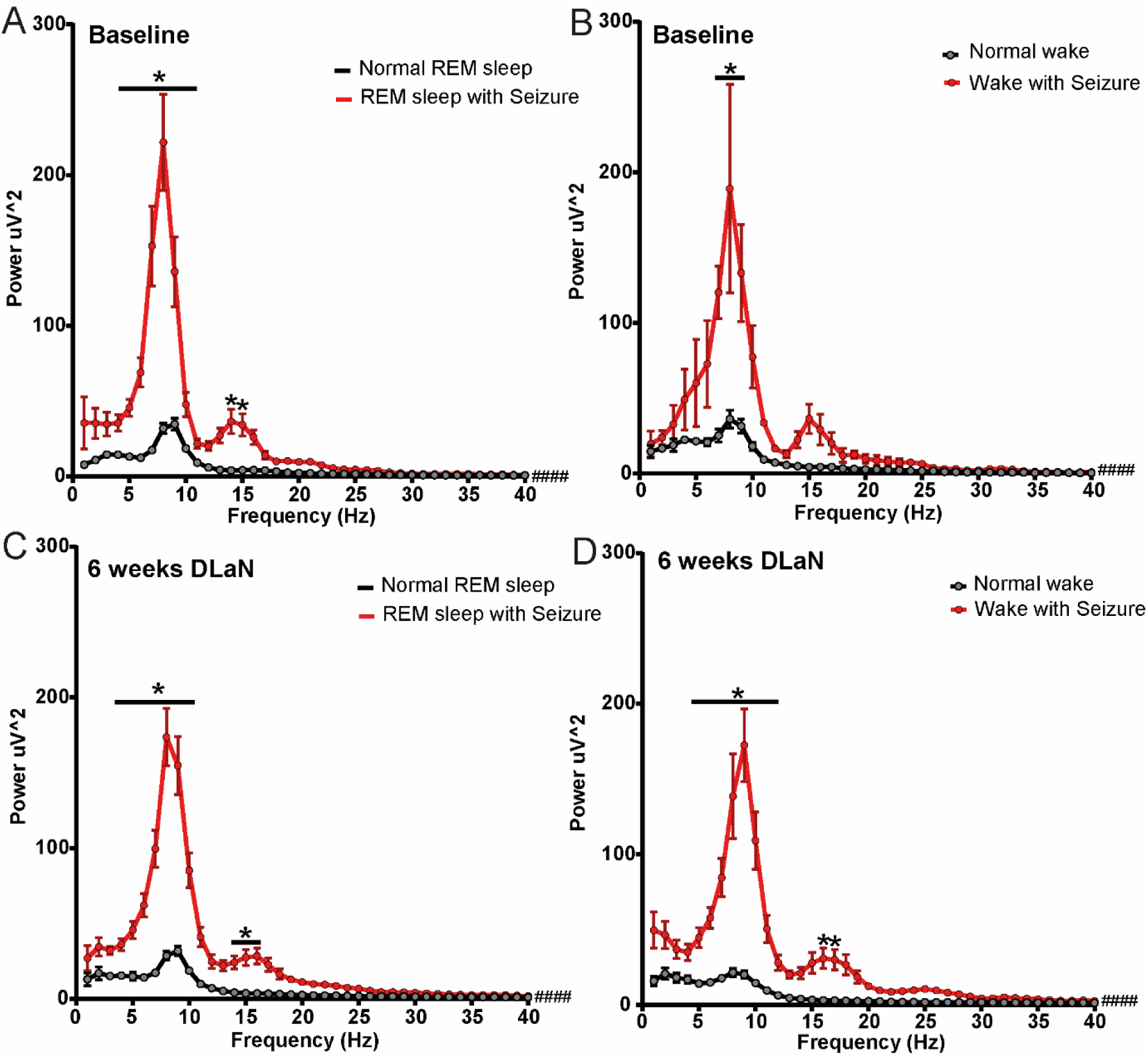
Seizure power analysis under baseline and 6 weeks of DLaN. Absolute EEG power spectrum during REM sleep (**A, C**) and waking (**B, D**) with seizure occurrence (red) and without seizure occurrence (black) for baseline and 6 weeks DLaN in *Cntnap2* KO mice. Pound sign indicates a significant interaction between the two factors “Frequency” and “DLaN” (#### p < 0.0001, two-way ANOVA). Asterisks indicate significant differences between abnormal EEG and normal EEG (*p < 0.05, **p < 0.01, Bonferroni multiple comparisons test after significant two-way ANOVA, interaction of “Frequency” and “seizure”). The frequency bin size is 1 Hz. n = 12–16 mice per condition. Data are shown as mean ± SEM.

